# Improved protein glycosylation enabled heterologous biosynthesis of monoterpenoid indole alkaloids and their unnatural derivatives in yeast

**DOI:** 10.1101/2022.06.08.495323

**Authors:** Mohammadamin Shahsavarani, Joseph Christian Utomo, Rahul Kumar, Melina Paz-Galeano, Jorge Jonathan Oswaldo Garza-García, Zhan Mai, Dae-Kyun Ro, Yang Qu

## Abstract

With over 3,000 reported structures, monoterpenoid indole alkaloids (MIAs) constitute one of the largest alkaloid groups in nature, including the clinically important anticancer drug vinblastine and its semi-synthetic derivatives from *Catharanthus roseus* (Madagascar’s periwinkle). With the elucidation of the complete 28-step biosynthesis for anhydrovinblastine, it is possible to investigate the heterologous production of vinblastine and other medicinal MIAs. In this study, we successfully expressed the flavoenzyme *O*-acetylstemmadenine oxidase in *Saccharomyces cerevisiae* (baker’s yeast) by signal peptide modification, which is a vinblastine biosynthetic gene that has not been functionally expressed in this system. We also report the simultaneous genomic integration of ∼18 kb MIA biosynthetic gene cassettes as single copies by CRISPR-Cas9 in baker’s yeast, which enabled the biosynthesis of vinblastine precursors catharanthine and tabersonine from the feedstocks secologanin and tryptamine. We further demonstrated the biosynthesis of fluorinated and hydroxylated catharanthine and tabersonine derivatives using our yeasts, which showed that the MIA biosynthesis accommodates unnatural substrates, and the system can be further explored to produce other complex MIAs.

With over 3,000 members, monoterpenoid indole alkaloids (MIA) are one of the largest and most diverse alkaloids in nature including many human medicines, such as chemotherapeutics vinblastine from *Catharanthus roseus* (Madagascar’s periwinkle) and camptothecin from *Camptotheca accuminata* (happy tree), and antiarrhythmic ajmaline from *Rauwolfia serpentina* (Indian snakeroot).^1^ Recent studies have elucidated the complete 28-step biosynthetic pathway for anhydrovinblastine in *C. roseus*, which involves diverting a primary monoterpene geranyl pyrophosphate into the biosynthesis of secologanin via the iridoid pathway (9 steps), genesis of the first MIA strictosidine that is the universal precursor to almost all MIAs (2 steps), conversion of strictosidine to iboga type MIA catharanthine and aspidosperma type tabersonine (9 steps), decorating tabersonine to vindoline (7 steps), and the final step that couples vindoline and catharanthine to make anhydrovinblastine (Fig. 1). ^2-12^ These studies not only revealed the remarkable complexity of MIA formations but also enabled the exploration in heterologous production of bioactive MIAs and intermediates that are usually found in low quantities in their natural sources. Notably, strictosidine and a related corynanthe type MIA ajmalicine have been produced *de novo* in *Saccharomyces cerevisiae* (baker’s yeast), ^13,14^ while vindoline has been produced in baker’s yeast from tabersonine feedstock. ^3,15,16^ For strictosidine production in yeast, the challenges lie in the generally low monoterpene biosynthesis output and the intermediates consumption by yeast native metabolism.^13,14,17^ While studies did not report rapid MIA consumption by yeast, vindoline yields were improved by optimizing the stoichiometry of cytochrome P450 monooxygenase (CYP), CYP redox partner CYP reductase (CPR), and other factors related with CYP activities such as endoplasmic reticulum (ER) homeostasis and NADPH co-factor regeneration that are commonly exploited.^15,16^ In this study, we constructed yeast strains containing the remaining vinblastine biosynthetic segment and produced catharanthine and tabersonine by feeding precursors, secologanin and tryptamine, as well as their unnatural derivatives by feeding substituted tryptamine.

## Results and Discussion

Previously, we have shown the production of a key MIA intermediate stemmadenine by feeding 19*E*-geissoschizine substrate, which was produced *in vitro* by geissoschizine synthase (GS), to baker’s yeast expressing the CYP geissoschizine oxidase (GO) and two reductases Redox 1 and 2 (strain 0).^5^ In this study, we further included the stemmadenine *O*-acetyltransferase (SAT), *O*-acetylstemmadenine oxidase (ASO), dihydroprecondylocarpine acetate synthase (DPAS), hydrolase 1 (HL1, catharanthine synthase), and hydrolase 2 (HL2, tabersonine synthase) in yeast to investigate the biosynthesis of catharanthine and tabersonine in this system (Fig. 1). ASO is a flavoprotein oxygenase and a member of the enzyme class Berberine Bridge Enzyme (BBE)-like proteins, which require eukaryotic glycosylation pathway to fold and mature into functional proteins.^2,18^ In addition to BBE itself involved in the benzylisoquinoline alkaloid biosynthesis from *Papaver somniferum* (opium poppy), another notable member is the tetrahydrocannabinolic acid synthase (THCAS) from *Cannabis sativa*. ^19^ These proteins contain a *N*-terminal signal peptide (SP) to direct the nascent polypeptide through ER lumen and trans-Golgi network for protein folding and *N*-glycosylation. ^20^ In cannabis, THCAS is secreted out of the cells similar to many other apoplastic glycoproteins, whereas BBE and ASO are re-routed to vacuoles and small vesicles, which is similar to the vacuolar glycoprotein carboxypeptidase Y (CPY) and proteinase A in baker’s yeast. ^7,21,22^ Previously we have shown that ASO, as either intact proteins with its own SP or *N*-terminal truncated proteins after removing the SP, could not be expressed to detectable levels in baker’s yeast, while active proteins could be purified by transient expression in *Nicotiana benthamiana* (tobacco) leaves. ^2^ The results were consistent with another report on unsuccessful ASO expression in baker’s yeast and other reports on the difficulty to produce functional BBE-like oxygenases without *N*-terminal SP modification in this system. ^7,18,19^ Recently, THCAS and a serine carboxypeptidase-like (SCPL) acyltransferase littorine synthase involved in plant tropane alkaloid biosynthesis that also require *N*-glycosylation were expressed functionally in baker’s yeast by swapping their native SP with those from yeast CPY or proteinase A. ^23,24^ The results suggest that plant SPs may not be effectively recognized by the *N*- glycosylation pathway in baker’s yeast and SP replacement may lead to functional expression of ASO in yeast.

**Figure 1.**
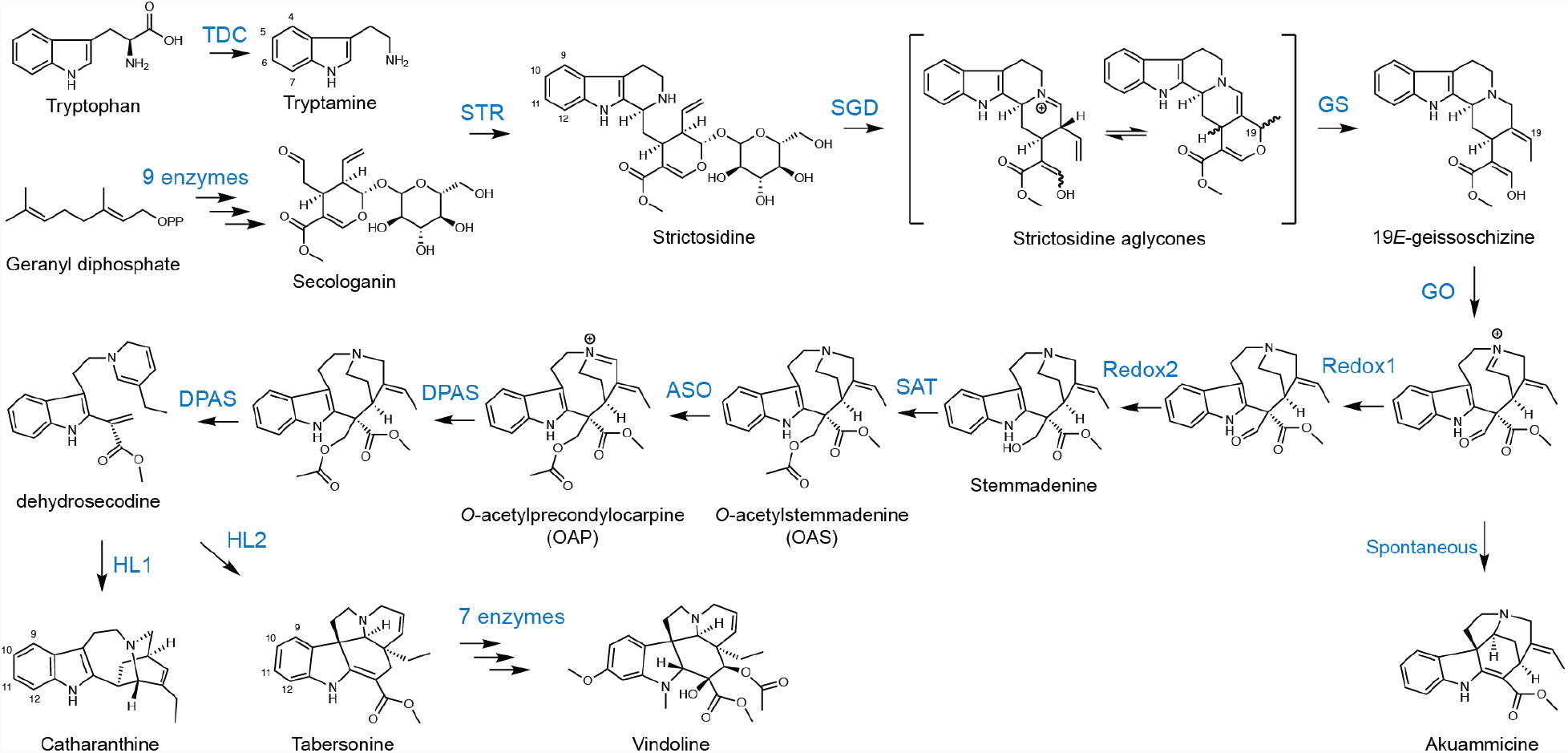
The biosynthetic pathway for monoterpenoid indole alkaloids (MIAs) catharanthine and vindoline in *C. roseus*, which couple to form the anticancer anhydrovinblastine and other derivatives. TDC: tryptophan decarboxylase (Genbank P17770); STR: strictosidine synthase (Genbank CAA43936); SGD: strictosidine β-glucosidase (Genbank AAF28800); GS: geissoschizine synthase (Genbank MF770507); GO: geissoschizine oxidase (Genbank MF770508); Redox1/2: oxidized geissoschizine reductase 1/2 (Genbank MF770509, MF770510); SAT: stemmadenine *O*-acetyltransferase (Genbank MF770511); ASO: *O*-acetylstemmadenine oxidase (Genbank MH136588); DPAS: dihydroprecondylocarpine synthase (Genbank A0A1B1FHP3); HL1: hydrolase 1/catharanthine synthase (Genbank MF770512); HL2: hydrolase 2/tabersonine synthase (Genbank MF770513).

We therefore generated two codon-optimized ASO with the addition of yeast CPY SP (amino acid AA1-34, Fig. 2, Supplementary Fig. 1). The first version (CPY-ASO) included the yeast CPY SP addition directly to the *N*-terminus of intact ASO, which resulted in two tandem *N*-terminal SPs. The second version (CPY-dASO) instead swapped ASO SP (AA1-24) with CPY SP. Both versions were expressed as *C*-terminal myc-tagged proteins under the control of Gal1 promoter in high-copy pESC-Ura vector (2μ origin of replication). The western blot showed successful protein expression and cleavage of SPs, since both versions of ASO were detected as approx. 55 kDa proteins while the expected size for CPY-ASO prior to SP removal is 63 kDa (Fig. 2).

**Figure 2.**
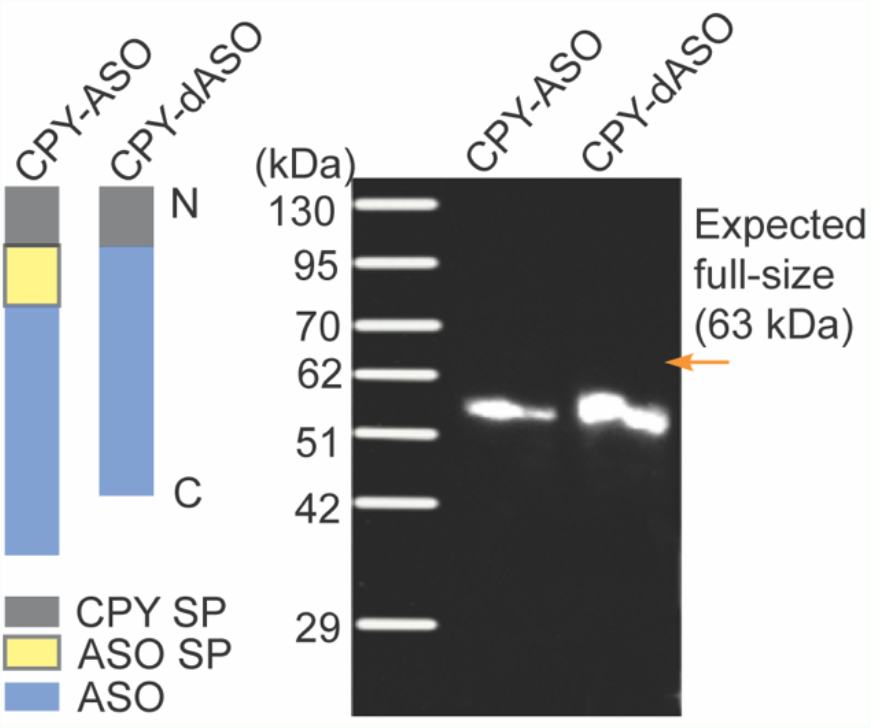
Placing yeast carboxypeptidase Y (CPY) signal peptide (SP) at *N*-terminus of *O*-acetylstemmadenine oxidase (ASO) allowed detectable expression of this flavoprotein oxygenase by western blot. Two codon-optimized ASO were synthesized. The CPY-ASO version contained a *N*-terminal CPY SP and ASO own SP, whereas the CPY-dASO version instead has ASO SP replaced by CPY SP (left panel). When expressed in yeast, ASO expression from both versions was detected by Western blots using anti-myc antibodies targeting the myc epitope tagged at *C*-termini of ASO (right panel). The reduced protein sizes (∼55 kDa) suggested successful removal of SP in both versions as the expected full-length protein was 63 kDa.

Next, we co-expressed GO, Redox1, Redox2, SAT, CPY-dASO, DPAS, and HL1 (Strain 1) or HL2 (strain 2) encoded in 2μ plasmids with 3 distinct selection markers under inducible Gal1/10 promoters in yeast (Fig. 3). When fed with 19*E*-geissoschizine, the strains successfully produced catharanthine and tabersonine, respectively (Supplementary Fig. 2 and 3). The conversion rates from geissoschizine were 0.28% for catharanthine and 0.36% for tabersonine (Fig. 3). When we replaced CPY-dASO with non-modified ASO containing native SP (strain 3), the catharanthine levels after geissoschizine feeding dropped by 71.9-fold to only 0.004% (Fig. 3). This result confirmed that SP swapping is necessary for functional expression of ASO in baker’s yeast.

**Figure 3.**
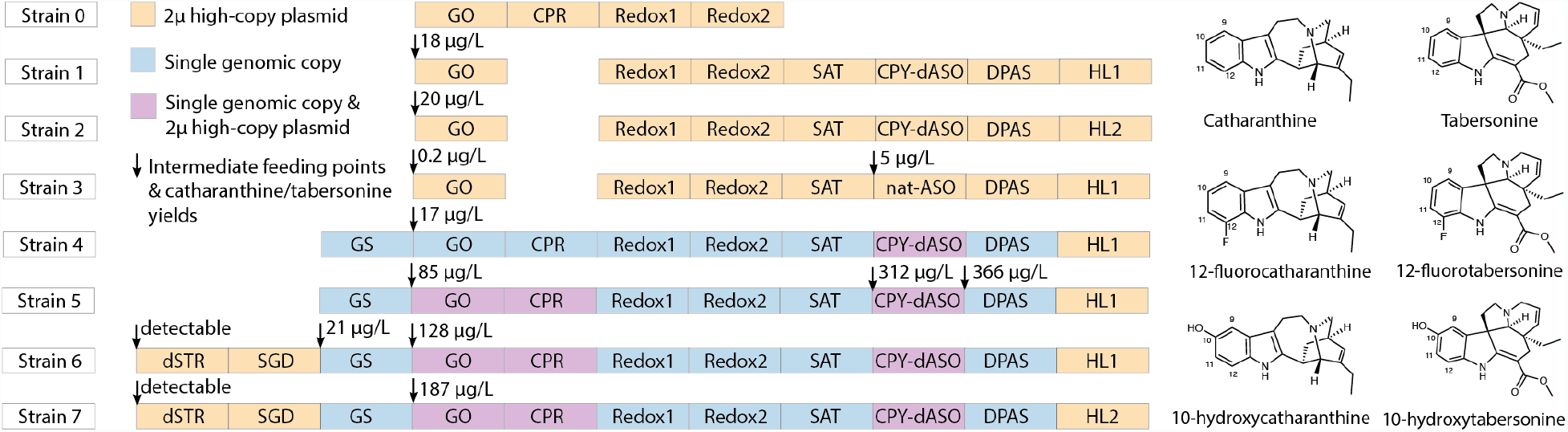
Various yeast strains with different alkaloid biosynthetic genes used in this study. The biosynthetic genes are shown in coloured blocks. Yellow colour indicates the expression from high-copy 2μ plasmids; blue colour indicates the expression from single genomic copies; and lavender colour indicates the expression from both high-copy 2μ plasmids and single genomic copies. All genes are under the control of Gal1/10 bi-directional promoters and their expression was induced by galactose in yeast media. The arrows show the points where pathway intermediates were fed to the yeasts, and the yields of catharanthine with HL1 activity and those of tabersonine with HL2 activities were indicated next to the arrow signs. By feeding yeasts with 12-fluorostrictosidine aglycones and 10-hydroxystrictosidine aglycones, respective substituted catharanthine and tabersonine were detected (right panel).

We further compared MIA biosynthesis in yeast between using high-copy plasmids and single genomic gene copies. A CRISPR-Cas9-compatible yeast strain was previously engineered to include identical gRNA-binding sequences that are placed adjacent to six high-expressing yeast genes, allowing simultaneous integration of up to six DNA cassettes as single copies in a single transformation. ^25^ Using this in-house strain, we integrated the genes GS, GO, CPR, Redox1, Redox2, SAT, CPY-dASO, and DPAS in four yeast genomic loci as single copies (Supplementary Fig. 4). The total length of integrated foreign genes was ∼18 kb over four loci, which is one of the largest foreign DNA integrations by multiplex CRISPR/Cas9 in yeast. ^26^ All MIA biosynthetic genes were under the control of galactose-inducible, bi-directional Gal1/10 promoters to avoid potential cytotoxicity from heterologous gene expression. The terminal enzymes HL1 and HL2 were instead expressed from 2μ plasmids, which directed the biosynthesis to either catharanthine or tabersonine by simple plasmid transformation (Fig. 3).

When fed with 19*E*-geissoschizine, the genomic integrated yeast with HL1 on plasmid (strain 4) successfully produced catharanthine at amounts comparable to strain 1 (0.26% geissoschizine conversion, Fig. 3). We further transformed GO and CPR on 2μ plasmid into strain 4 to generate strain 5, which resulted in 4.9-fold increase (1.27% geissoschizine conversion, 85 μg L^-1^) in catharanthine production and suggested that the CYP GO activity was a limiting factor in strain 4. In strain 5 feeding experiment, 83.6% of 15 mg L^-1^ 19*E*-geissoschizine substrate was consumed by yeast. The only other detectable MIA was a pathway by-product akuammicine, decomposed from GO reaction, comprising 9.6% of substate (Supplementary Fig. 5).

Since only small fraction of geissoschizine was converted to catharanthine, we investigated strain 5 using downstream intermediates *O*-acetylstemmadenine (OAS, product of SAT) and *O*-acetylprecondylocarpine (OAP, product of ASO, Fig. 1). These two substrates are the few stable intermediate in this section of pathway as many other intermediates rapidly decompose, which is likely a major limitation for MIA biosynthesis in heterologous system. When we fed OAS (7.5 mg L^-1^) to strain 5 overnight, 91.0% of substrate was consumed by yeast, and other detectable MIAs included catharanthine (11.0% substrate, 312 μg L^-1^), OAP (31.9% substrate), and the deacetylated stemmadenine (14.6% substrate, Fig. 3, Supplementary Fig. 5). When we fed OAP (6 mg L^-1^) to strain 5, 91.2% substrate was consumed, and other detectable MIAs included catharanthine (16.2% substrate, 366 μg L^-1^) and the deacetylated precondylocarpine (14.4% substrate, Fig. 3, Supplementary Fig. 5). These low conversion rates suggests that majority of the MIA intermediates were lost due to their instability in yeast environment.

To complete catharanthine and tabersonine biosynthesis in yeast from simple feedstocks tryptamine and secologanin, we included upstream genes strictosidine synthase (STR), strictosidine β-glucosidase (SGD) on a 2μ plasmid in strain 5 background to create strain 6 (HL1) and strain 7 (HL2). We opted to use a codon-optimized STR lacking its *N*-terminal vacuole targeting SP (AA1-18, Supplementary fig. 1) to localize STR to yeast cytosol, where other downstream enzymes reside. After secologanin and tryptamine substrate feeding, strain 6 produced detectable levels of catharanthine (Fig. 3, Supplementary fig. 6). Feeding downstream intermediates strictosidine aglycones (15 mg L^-1^, product of SGD) and 19*E*-geissoschizine (15 mg L^-1^) produced from *in vitro* reactions (Supplementary fig. 2), we observed 230-fold increase in catharanthine production (21 μg L^-1^) and 1400-fold (128 μg L^-1^) with respective substrates (Fig. 3, Supplementary fig. 6), suggesting poor efficiency of STR and SGD in living yeast cells. When using strain 7 with HL2, we obtained similar results for tabersonine production (Fig. 3, Supplementary fig. 6).

STR and SGD have been reported to accept unnatural substrates containing indole substitutions such as fluorine and methyl groups. ^27,28^ The enzymes downstream of SGD also accepted indole chlorinated substrates when tested in *C. roseus* hairy root cultures. ^29^ We therefore tested MIA biosynthesis with 7-fluorotryptamine, 5-chlorotryptamine, and 5-hydroxytryptamine (serotonin). Using recombinant enzymes purified from *E. coli*, STR, SGD, and GS successfully accepted both 7-fluorotryptamine and serotonin, and produced 12-flouro-and 10-hydroxygeissoschizine, while 5-chlorotryptamine was not accepted by STR (Supplementary Fig. 2). Feeding 7-fluorotryptamine or serotonine to strain 6 and 7 did not lead to detectable MIA accumulation, likely due to the low overall conversion rates in these strains and reduced enzyme activities with unnatural substrates. However, feeding 12-fluorostrictosidine aglycones and 10-hydroxystrictosidine aglycones to strain 6 and 7 led to detectable accumulations of respectively substituted catharanthine and tabersonine (Fig. 3, Supplementary fig. 6), indicating that the entire downstream enzymes can also accommodate unnatural substrates.

Previous studies have shown the *de novo* biosynthesis of strictosidine, and the vindoline biosynthesis from tabersonine in baker’s yeast. ^3,13-16^ In this study, we demonstrated the biosynthesis of catharanthine and tabersonine that are required for vinblastine biosynthesis from tryptamine, secologanin, and other downstream MIA intermediates. While the complete vinblastine biosynthesis in yeast requires significant pathway optimizations, our prior and current studies show that yeast can be used to produce unnatural and rare intermediates that may be used to explore the elusive biosynthesis of other medicinal MIAs.

## Materials and methods

### Yeast strains, cloning, and plasmid combinations

The yeast strain BY4742 (MATα, HIS3Δ1, LEU2Δ0, LYS2Δ0, URA3Δ0) were transformed with various alkaloid biosynthetic genes encoded on pESC-Leu, -His, -Ura vectors. The primers and gene cloning combinations are found in Supplementary table 1 and 2. Codon-optimized CPY-ASO and dSTR were synthesized and subcloned in pET-Duet (BamHI/HindIII) and pUG19 (NotI/ClaI) (Bio Basic Inc., Toronto, Canada) and their sequences are listed in Supplementary fig.1. The yeast strain SBY104 (MATα, HIS3Δ1, LEU2Δ0, LYS2Δ0, URA3Δ0) was used for CRISPR-Cas9 based genomic gene integration as described. ^25^ Briefly, eight MIA genes in four bi-directional cassettes were amplified using primers 31-40 and cloned into four donor DNA plasmids using Gibson Assembly method (Supplementary fig. 4). The genes are distributed as follows, iADH1 (H1):: GO-CPR; iPDC1 (H2):: GS-Redox 2, iPGK1 (H5):: DPAS-Redox 1, and iCDC19 (H7):: CPY-dASO-SAT. The plasmid combinations used to transform strain 0-6 is listed in Supplementary table 3.

### Multiplex CRISPR-Cas9-based integration

From the four donor plasmids, linear donor DNA fragments were amplified using repliQa HiFi ToughMix (Quantabio, Beverly, MA, USA) and primers 41-48. Yeast transformation was performed using the lithium acetate method as previously described. ^25^ Briefly, overnight yeast culture was diluted 100 times into 25 ml YPD media and incubated until OD_600_ reached 0.4 – 0.8. The cells were collected by centrifugation and washed with H_2_O. Approx. 2 × 10^7^ cells were incubated in 200 μl of 60% (w/v) PEG, 1 μg of pESC-URA::Cas9-gRNA plasmid, and 2 μg of each linear donor DNA for 15 min, which was followed by adding 50 μl of 2 mg/ml boiled salmon sperm DNA and 18 μl of 2 M lithium acetate into the mixtures. The mixtures were incubated at 42 °C for 40 min, immediately put on ice, spun down, and incubated in synthetic complex (SC)-Ura with glucose media for 2 hr at 30 °C before being plated on SC-Ura plates. The plates were incubated overnight at 37 °C and then incubated at 30 °C for 2-3 days. To confirm the genomic gene integrations, the yeast genomic DNA was extracted with Yeast DNA Extraction Kit (Thermo Fisher, Rockford, IL, USA). Three primers, which would produce an 800 bp product for negative colonies and a 2-3 kb product for positive colonies, were used to genotype each colony in each integration site. The primers used are: loci H1, primer 41, 49, 53; H2, 43, 50, 54; H5, 45,51,55; and H7, 47,52,56 (Supplementary table 1).

### Yeast cultivation

The yeast strains were grown in 1 ml SC drop-out media (Sigma Aldrich, St. Louis, MO, USA) with 2% (w/v) glucose in test tubes at 30 °C in a shaking incubator overnight, then the cells were collected by centrifugation, washed once with water, resuspended in 1 ml fresh media with 2% (w/v) galactose, and cultured in test tubes at 30 °C in a shaking incubator for 24 hr. The cells were collected by centrifugation and resuspended in 300 μL 20 mM pH 7.5 Tris-HCl buffer supplemented with various MIA substrates overnight. The culture supernatants were mixed with equal volume of methanol for LC-MS/MS analyses.

### LC-MS/MS and alkaloid identification

LC-MS/MS was performed on an Agilent Ultivo Triple Quadrupole LC-MS equipped with an Avantor® ACE® UltraCore™ SuperC18™ column (2.5 μm, 50×3mm), which included the solvent systems: solvent A, methanol: acetonitrile: ammonium acetate 1 M: water at 29:71:2:398; solvent B, methanol: acetonitrile: ammonium acetate 1 M: water at 130:320:0.25:49.7. The following linear gradient (8 min, 0.6 ml/min) were used: 0 min 80% A, 20% B; 0.5 min, 80% A, 20%B; 5.5 min 1% A, 99% B; 5.8 min 1% A, 99% B; 6.5 min 80% A, 20% B; 8 min 80% A, 20% B. The photodiode array detector records from 200 to 500 nm. The MS/MS was operated with gas temperature at 300°C, gas flow of 10 L/min, capillary voltage 4 kV, fragmentor 135 V, collision energy 30V with positive polarity. Catharanthine, tabersonine, and secologanin standards were purchased from Sigma-Aldrich Inc. (St. Louis, MO, USA). Trptamine, 7-fluorotryptamine, 5-chlorotryptamine, and serotonin were purchased from Cayman Chemicals (Ann Arbor, MI, USA). *O*-acetylstemmadenine and *O*-acetylprecondylocarpine were purified from a *C. roseus* mutant described previously. ^2^ The remaining intermediates/standards were produced, purified, and identified by NMR as described. ^4,5^

## References

1. Facchini, P. J. & De Luca, V. (2008) Opium poppy and Madagascar periwinkle: model non-model systems to investigate alkaloid biosynthesis in plants. The Plant Journal 54, 763–784.

2. Qu, Y., Safonova, O. & De Luca, V. (2018) Completion of the canonical pathway for assembly of anticancer drugs vincristine/vinblastine in Catharanthus roseus. The Plant journal 97, 257–266.

3. Qu, Y. et al. (2015) Completion of the seven-step pathway from tabersonine to the anticancer drug precursor vindoline and its assembly in yeast. Proceedings of the National Academy of Sciences 112, 6224–6229.

4. Qu, Y. et al. (2017) Geissoschizine synthase controls flux in the formation of monoterpenoid indole alkaloids in a Catharanthus roseus mutant. Planta 25, 1–10.

5. Qu, Y. et al. (2018) Solution of the multistep pathway for assembly of corynanthean, strychnos, iboga, and aspidosperma monoterpenoid indole alkaloids from 19 E-geissoschizine. Proceedings of the National Academy of Sciences 21, 201719979–6.

6. Geu-Flores, F., Sherden, N.H., Courdavault, V., Burlat, V., Glenn, W.S., Wu, C., Nims, E., Cui, Y. and O’Connor, S.E. (2012) An alternative route to cyclic terpenes by reductive cyclization in iridoid biosynthesis. Nature 492, 138–142.

7. Caputi, L. et al. (2018) Missing enzymes in the biosynthesis of the anticancer drug vinblastine in Madagascar periwinkle. Science 360, 1235–1239.

8. Sottomayor, M. & Barceló, A. R. (2003) Peroxidase from Catharanthus roseus (L.) G. Don and the biosynthesis of alpha-3’,4’-anhydrovinblastine: a specific role for a multifunctional enzyme. Protoplasma 222, 97–105.

9. Asada, K., Salim, V., Masada-Atsumi, S., Edmunds, E., Nagatoshi, M., Terasaka, K., Mizukami, H., De Luca, V. (2013) A 7-deoxyloganetic acid glucosyltransferase contributes a key step in secologanin biosynthesis in Madagascar periwinkle. The Plant Cell 25, 4123–4134

10. Salim, V., Wiens, B., Masada-Atsumi, S., Yu, F., De Luca, V. (2014) 7-Deoxyloganetic acid synthase catalyzes a key 3 step oxidation to form 7-deoxyloganetic acid in Catharanthus roseus iridoid biosynthesis. Phytochemistry 101, 23–31

11. Salim V., Yu, F., Altarejos., J., De Luca, V. (2013) Virus-induced gene silencing identifies Catharanthus roseus 7-deoxyloganic acid-7-hydroxylase, a step in iridoid and monoterpene indole alkaloid biosynthesis. The Plant Journal 76, 754–765

12. Miettinen, K., Dong, L., Navrot, N. et al. (2014) The seco-iridoid pathway from Catharanthus roseus. Nature Communication. 5, 3606 and erratum (2014) 5, 4175.

13. Brown, S., Clastre, M., Courdavault, V. & O’Connor, S. E. (2015) De novo production of the plant-derived alkaloid strictosidine in yeast. Proceedings of the National Academy of Sciences 112, 3205–3210.

14. Liu, T., Gou, Y., Zhang, B., Gao, R., Dong, C., Qi, M., Jiang, L., Ding, X., Li, C., Lian, J. (2022) Construction of ajmalicine and sanguinarine de novo biosynthetic pathways using stable integration sites in yeast. Biotechnology and Bioengineering 119, 1314–1326

15. Liu, T., Huang, Y., Jiang, L., Dong, C., Gou, Y., Lian, J. (2021) Efficient production of vindoline from tabersonine by metabolically engineered Saccharomyces cerevisiae. Communications Biology 4, 1089

16. Cruz, P. L. et al. (2021) Optimization of Tabersonine Methoxylation to Increase Vindoline Precursor Synthesis in Yeast Cell Factories. Molecules 26, 3596.

17. Campbell, A., Bauchart, P., Gold, N. D., Zhu, Y., De Luca, V., Martin, V. J. J. (2016) Engineering of a Nepetalactol-Producing Platform Strain of Saccharomyces cerevisiae for the Production of Plant Seco-Iridoids. ACS Synthetic Biology 5, 405–414

18. Winkler, A., Lyskowski, A., Riedl, S., Puhl, M., Kutchan, T. M., Macheroux, P., Gruber, K. (2008) A concerted mechanism for berberine bridge enzyme. Nature Chemical Biology 4, 739–741.

19. Sirikantaramas, S., Morimoto, S., Shoyama, Y., Ishikawa, Y., Wada, Y., Shoyama, Y., Taura, F. (2004) The gene controlling marijuana psychoactivity: molecular cloning and heterologous expression of Δ1-tetrahydrocannabinolic acid synthase from Cannabis sativa L. Journal of Biological Chemistry 279, 39767–39774.

20. Nagashima, Y., Schaewen von, A., Koiwa, H. (2018) Function of N-glycosylation in plants. Plant Science 274, 70–79.

21. Sirikantaramas, S., Taura, F., Tanaka, Y., Ishikawa, Y., Morimoto, S., Shoyama, Y. (2005) Tetrahydrocannabinolic Acid Synthase, the Enzyme Controlling Marijuana Psychoactivity, is Secreted into the Storage Cavity of the Glandular Trichomes. Plant Cell Physiology 46, 1578–1582.

22. Bird D. A., Facchini P. J. (2001) Berberine bridge enzyme, a key branch-point enzyme in benzylisoquinoline alkaloid biosynthesis, contains a vacuolar sorting determinant. Planta 213, 888–897

23. Luo, X., Reiter, M. A., Espaux, L. D. X., Wong, J., Denby, C. M., Lechner, A., Zhang, Y., Grzybowski, A. T., Harth, S., Lin, W., Lee, H., Yu, C., Shin, J., Deng, K., Benites, V. T., Wang, G., Baidoo, E. E. K., Chen, Y., Dev, I., Petzold, C. J., Keasling, J. D. (2019) Complete biosynthesis of cannabinoids and their unnatural analogues in yeast. Nature 567, 123–126.

24. Srinivasan, P., Smolke C. D. (2020) Biosynthesis of medicinal tropane alkaloids in yeast. Nature 585, 614–619

25. Baek, S., Utomo, J. C., Lee, J. Y., Dalal, K., Yoon, Y. J., and Ro, D. K. (2021). The yeast platform engineered for synthetic gRNA-landing pads enables multiple gene integrations by a single gRNA/Cas9 system. Metabolic Engineering 64, 111–121

26. Utomo J. C., Hodgins, C. L., Ro, D. K. (2021) Multiplex Genome Editing in Yeast by CRISPR/Cas9 – A Potent and Agile Tool to Reconstruct Complex Metabolic Pathways. Frontiers in Plant Science 12, 719148 http://doi.org/10.3389/fpls.2021.719148

27. McCoy, E., Galan, M. C., O’Connor, S. E. (2006) Substrate specificity of strictosidine synthase. Bioorganic & Medicinal Chemistry Letters 16, 2475–2478.

28. Yerkes, N., Wu, J., McCoy, E., Galan, M. C., Chen, S., O’Connor, S. E. (2008) Substrate Specificity and Diastereoselectivity of Strictosidine Glucosidase, a Key Enzyme in Monoterpene Indole Alkaloid Biosynthesis. Bioorganic & Medicinal Chemistry Letters 18, 3095–3098.

29. Runguphan, W., Qu, X., O’Connor, S. E. (2010) Integrating carbon–halogen bond formation into medicinal plant metabolism. Nature 468, 461–464

