## Supplementary information for "Improved protein glycosylation enabled heterologous biosynthesis of monoterpenoid indole alkaloids and their unnatural derivatives in yeast"

**A**

CPY-ASO

MKAFTSLLCGLGLSTTLAKAISLQRPLGLDKDVLELMIKKVPIVLSIFCFLLLLSSSHGSELIPEAFLNCISNKFSLDVSILNILHVPSNSSYDSVLKSTI
QNPRFLKSPKPLAIITPVLHSHVQSAVICTKQAGLQIRIRSGGADYEGLSYRSEVPFILLDLQNLRSISVDIEDNSAWVESGATIGEFYHEIAQNSPVH
AFPAGVSSSVGIGGHLSSGGFGTLLRKYGLAADNIIDAKIVDARGRILDRESMGEDLFWAIRGGGGASFGVIVSWKVKLVKVPPMVTVFILSKTYE
EGGLDLLHKWQYIEHKLPEDLFLAVSIMDDSSSGNKTLMAGFMSLFLGKTEDLLKVMAENFPQLGLKKEDCLEMNWIDAAMYFSGHPIGESRSV
LKNRESHLPKTCVSIKSDFIQEPQSMDALEKLWKFCREEENSPIILMLPLGGMMSKISESEIPFPYRKDVIYSMIYEIVWNCEDDESSEEYIDGLGRL
EELMTPYVKQPRGSWFSTRNLYTGKNKGPGTTYSKAKEWGFRYFNNNFKKLALIKGQVDPENFFYYEQSIPPLHLQVEL

CPY-dASO

MKAFTSLLCGLGLSTTLAKAISLQRPLGLDKDVLELIPEAFLNCISNKFSLDVSILNILHVPSNSSYDSVLKSTIQNPRFLKSPKPLAIITPVLHSHVQS
AVICTKQAGLQIRIRSGGADYEGLSYRSEVPFILLDLQNLRSISVDIEDNSAWVESGATIGEFYHEIAQNSPVHAFPAGVSSSVGIGGHLSSGGFGTL
LRKYGLAADNIIDAKIVDARGRILDRESMGEDLFWAIRGGGGASFGVIVSWKVKLVKVPPMVTVFILSKTYEEGGLDLLHKWQYIEHKLPEDLFL
AVSIMDDSSSGNKTLMAGFMSLFLGKTEDLLKVMAENFPQLGLKKEDCLEMNWIDAAMYFSGHPIGESRSVLKNRESHLPKTCVSIKSDFIQEPQ
SMDALEKLWKFCREEENSPIILMLPLGGMMSKISESEIPFPYRKDVIYSMIYEIVWNCEDDESSEEYIDGLGRLEELMTPYVKQPRGSWFSTRNLYT
GKNKGPGTTYSKAKEWGFRYFNNNFKKLALIKGQVDPENFFYYEQSIPPLHLQVEL

**B**

CPY-ASO

ATGAAAGCATTCACCAGTTTACTATGTGGACTAGGCCTGTCCACTACACTCGCTAAGGCCATCTCATTGCAAAGACCGTTGGGTCTAGATAAGGACGTTTTGGAGCTCATGATAAAAAA
AGTCCCAATAGTTCTTTCAATTTTCTGCTTTCTTCTTCTACTCTCATCATCCCATGGCTCAGAGCTCATTCCTGAAGCTTTTCTCAATTGTATTTCCAATAAATTTTCATTAGATGTAT
CCATTTTAAACATTCTTCATGTTCCCAGCAATTCTTCCTATGATTCTGTTCTCAAATCTACTATCCAAAATCCAAGATTCCTCAAATCACCCAAGCCCTTAGCTATAATCACCCCAGTA
CTTCACTCCCATGTCCAATCTGCTGTTATCTGTACCAAACAAGCCGGTTTACAAATTAGAATCCGAAGCGGAGGAGCTGATTACGAGGGCTTATCCTATCGTTCTGAGGTTCCCTTTAT
TCTGCTAGATCTCCAGAATCTTCGATCAATTTCCGTTGATATTGAAGACAACAGCGCTTGGGTCGAATCAGGAGCAACAATTGGTGAATTCTATCATGAGATAGCTCAGAACAGCCCTG
TTCATGCGTTTCCAGCTGGGGTCTCTTCCTCTGTTGGAATTGGCGGCCATTTGAGTAGCGGCGGTTTTGGTACATTGCTTCGGAAATATGGATTAGCAGCCGATAATATAATCGATGCA
AAAATTGTTGATGCCAGAGGCAGAATTCTTGATAGGGAATCAATGGGAGAAGATCTATTTTGGGCTATTAGAGGAGGAGGAGGAGCTAGTTTTGGTGTTATAGTTTCTTGGAAGGTTAA
ACTTGTAAAAGTCCCTCCGATGGTAACTGTTTTCATCTTGTCCAAGACTTATGAAGAAGGAGGTTTAGATCTTCTACACAAATGGCAATATATAGAACACAAACTCCCTGAAGATTTAT
TCCTTGCTGTAAGCATCATGGATGATTCATCTAGTGGAAATAAAACACTTATGGCAGGTTTTATGTCTCTGTTTCTTGGAAAAACAGAGGACCTTCTGAAAGTAATGGCGGAAAATTTC
CCACAACTTGGATTGAAAAAGGAAGATTGCTTAGAAATGAATTGGATTGATGCAGCAATGTATTTTTCAGGACACCCAATTGGAGAATCCCGATCTGTGCTTAAAAACCGAGAATCTCA
TCTTCCAAAGACATGCGTTTCGATCAAATCAGACTTTATTCAAGAACCACAATCCATGGATGCATTGGAAAAGTTATGGAAGTTTTGTAGGGAAGAAGAAAATAGTCCCATAATACTGA
TGCTTCCACTGGGGGGAATGATGAGTAAAATATCAGAATCAGAAATCCCATTTCCTTACAGAAAAGATGTGATTTACAGTATGATATACGAAATAGTTTGGAATTGTGAAGACGATGAA
TCATCGGAAGAATATATCGATGGATTGGGAAGGCTTGAGGAATTAATGACTCCATATGTGAAACAACCAAGAGGTTCTTGGTTCAGCACCAGAAACCTTTATACCGGTAAAAATAAAGG
TCCAGGAACAACTTATTCCAAAGCTAAAGAATGGGGATTTCGGTATTTTAATAATAATTTCAAAAAGTTGGCCCTTATCAAAGGACAAGTTGATCCAGAAAACTTCTTCTACTATGAAC
AAAGCATTCCCCCTCTCCATTTACAAGTCGAACTT

CPY-dASO

ATGAAAGCATTCACCAGTTTACTATGTGGACTAGGCCTGTCCACTACACTCGCTAAGGCCATCTCATTGCAAAGACCGTTGGGTCTAGATAAGGACGTTTTGGAGCTCATTCCTGAAGC
TTTTCTCAATTGTATTTCCAATAAATTTTCATTAGATGTATCCATTTTAAACATTCTTCATGTTCCCAGCAATTCTTCCTATGATTCTGTTCTCAAATCTACTATCCAAAATCCAAGAT
TCCTCAAATCACCCAAGCCCTTAGCTATAATCACCCCAGTACTTCACTCCCATGTCCAATCTGCTGTTATCTGTACCAAACAAGCCGGTTTACAAATTAGAATCCGAAGCGGAGGAGCT
GATTACGAGGGCTTATCCTATCGTTCTGAGGTTCCCTTTATTCTGCTAGATCTCCAGAATCTTCGATCAATTTCCGTTGATATTGAAGACAACAGCGCTTGGGTCGAATCAGGAGCAAC
AATTGGTGAATTCTATCATGAGATAGCTCAGAACAGCCCTGTTCATGCGTTTCCAGCTGGGGTCTCTTCCTCTGTTGGAATTGGCGGCCATTTGAGTAGCGGCGGTTTTGGTACATTGC
TTCGGAAATATGGATTAGCAGCCGATAATATAATCGATGCAAAAATTGTTGATGCCAGAGGCAGAATTCTTGATAGGGAATCAATGGGAGAAGATCTATTTTGGGCTATTAGAGGAGGA
GGAGGAGCTAGTTTTGGTGTTATAGTTTCTTGGAAGGTTAAACTTGTAAAAGTCCCTCCGATGGTAACTGTTTTCATCTTGTCCAAGACTTATGAAGAAGGAGGTTTAGATCTTCTACA
CAAATGGCAATATATAGAACACAAACTCCCTGAAGATTTATTCCTTGCTGTAAGCATCATGGATGATTCATCTAGTGGAAATAAAACACTTATGGCAGGTTTTATGTCTCTGTTTCTTG
GAAAAACAGAGGACCTTCTGAAAGTAATGGCGGAAAATTTCCCACAACTTGGATTGAAAAAGGAAGATTGCTTAGAAATGAATTGGATTGATGCAGCAATGTATTTTTCAGGACACCCA
ATTGGAGAATCCCGATCTGTGCTTAAAAACCGAGAATCTCATCTTCCAAAGACATGCGTTTCGATCAAATCAGACTTTATTCAAGAACCACAATCCATGGATGCATTGGAAAAGTTATG
GAAGTTTTGTAGGGAAGAAGAAAATAGTCCCATAATACTGATGCTTCCACTGGGGGGAATGATGAGTAAAATATCAGAATCAGAAATCCCATTTCCTTACAGAAAAGATGTGATTTACA
GTATGATATACGAAATAGTTTGGAATTGTGAAGACGATGAATCATCGGAAGAATATATCGATGGATTGGGAAGGCTTGAGGAATTAATGACTCCATATGTGAAACAACCAAGAGGTTCT
TGGTTCAGCACCAGAAACCTTTATACCGGTAAAAATAAAGGTCCAGGAACAACTTATTCCAAAGCTAAAGAATGGGGATTTCGGTATTTTAATAATAATTTCAAAAAGTTGGCCCTTAT
CAAAGGACAAGTTGATCCAGAAAACTTCTTCTACTATGAACAAAGCATTCCCCCTCTCCATTTACAAGTCGAACTT

**C**

dSTR

ATGAGTTCTAGTTCACCAATTTTAAAAAAGATTTTTATTGAATCTCCATCTTATGCTCCAAATGCTTTTACATTTGATTCTACTGATAAAGGTTTTTATACTTCTGTTCAAGATGGTAG
AGTTATTAAGTATGAAGGTCCTAATTCTGGTTTTACTGATTTTGCATACGCTTCTCCATTTTGGAATAAGGCTTTTTGTGAAAATTCAACTGATCCTGAAAAAAGACCTTTGTGTGGTA
GAACATACGATATTTCTTATGATTATAAGAATAGTCAAATGTACATTGTTGATGGTCATTATCATTTGTGTGTTGTTGGTAAAGAAGGTGGTTATGCTACACAACTAGCAACTTCTGTT
CAAGGTGTTCCTTTTAAATGGTTATACGCTGTTACAGTAGATCAAAGAACTGGTATTGTTTATTTTACAGATGTTTCATCAATTCATGATGATTCACCTGAAGGTGTTGAAGAAATTAT
GAATACATCTGATAGAACTGGTAGATTGATGAAATATGATCCATCTACTAAAGAAACAACATTATTGTTGAAAGAATTACATGTTCCAGGTGGTGCTGAAATTTCAGCTGATGGTTCTT
TTGTTGTTGTTGCAGAATTTTTATCTAATAGAATTGTTAAATATTGGTTAGAAGGTCCTAAAAAGGGTTCAGCTGAATTTTTAGTTACTATTCCTAATCCAGGTAATATCAAAAGAAAT
TCTGATGGTCATTTTTGGGTTTCTTCATCTGAAGAATTAGATGGTGGTCAACATGGTAGAGTTGTTAGTAGAGGTATTAAGTTTGATGGTTTTGGTAATATTTTGCAAGTTATCCCATT
GCCACCACCATACGAAGGTGAACATTTCGAACAAATTCAAGAACACGACGGTTTGTTGTACATTGGTTCCTTGTTCCACTCCTCAGTCGGTATCTTGGTCTACGACGACCATGATAACA
AGGGTAACTCCTACGTCAGCTCC

**Supplementary figure 1. The nucleotide sequences (A) and amino acid sequences (B) of codon-optimized *O*-acetylstemmadenine oxidase (ASO) with both yeast carboxypeptidase Y (CPY) signal peptide (SP) and ASO own SP (CPY-ASO), and a single CPY SP without ASO own SP (CPY-dASO), and nucleotide sequences (C) of codon-optimized dSTR.** Both ASO versions were subcloned in pESC-Ura vector within BamHI/SalI sites. The CPY SP (amino acid 1-34) is shaded in grey, whereas the ASO own SP is highlighted in yellow. CPY-dASO was created by cleaving ASO own SP between the SacI sites (magenta highlight).

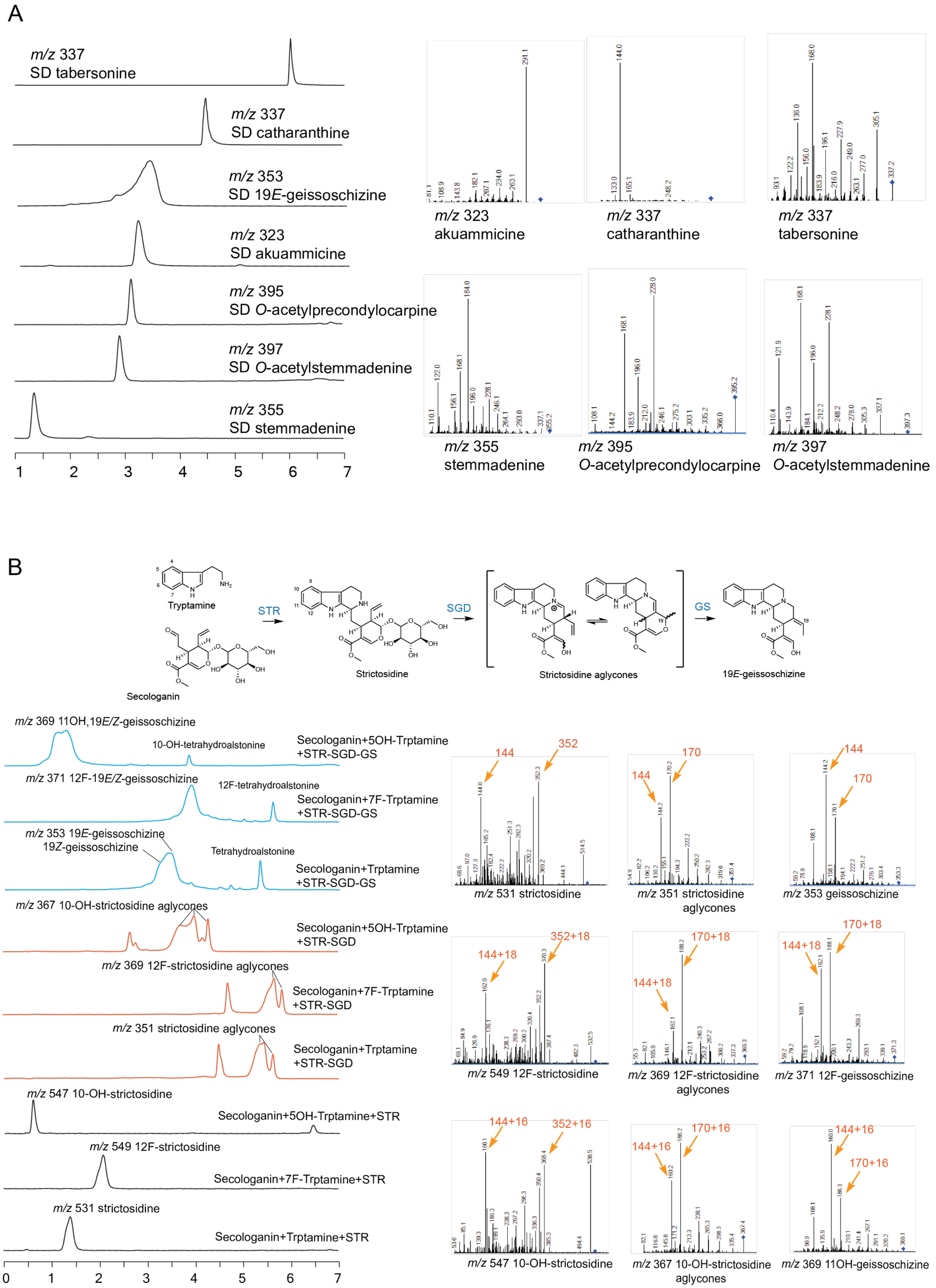

A
m/z 337
SD tabersonine
m/z 337
SD catharanthine
m/z 353
SD 19E-geissoschizine
m/z 323
SD akuammicine
m/z 395
SD O-acetylprecondylocarpine
m/z 397
SD O-acetylstemmadenine
m/z 355
SD stemmadenine
1 2 3 4 5 6 7
m/z 323
akuammicine
m/z 337
catharanthine
m/z 337
tabersonine
m/z 355
stemmadenine
m/z 395
O-acetylprecondylocarpine
m/z 397
O-acetylstemmadenine
B
Tryptamine
STR
SGD
GS
Strictosidine
Strictosidine aglycones
19E-geissoschizine
Secologanin
m/z 369 11OH,19E/Z-geissoschizine
10-OH-tetrahydroalstonine
Secologanin+5OH-Trptamine
+STR-SGD-GS
m/z 371 12F-19E/Z-geissoschizine
12F-tetrahydroalstonine
Secologanin+7F-Trptamine
+STR-SGD-GS
m/z 353 19E-geissoschizine
19Z-geissoschizine
Tetrahydroalstonine
Secologanin+Trptamine
+STR-SGD-GS
m/z 367 10-OH-strictosidine aglycones
Secologanin+5OH-Trptamine
+STR-SGD
m/z 369 12F-strictosidine aglycones
Secologanin+7F-Trptamine
+STR-SGD
m/z 351 strictosidine aglycones
Secologanin+Trptamine
+STR-SGD
m/z 547 10-OH-strictosidine
Secologanin+5OH-Trptamine+STR
m/z 549 12F-strictosidine
Secologanin+7F-Trptamine+STR
m/z 531 strictosidine
Secologanin+Trptamine+STR
0 1 2 3 4 5 6 7
144
352
m/z 531 strictosidine
144
170
m/z 351 strictosidine
aglycones
144
170
m/z 353 geissoschizine
144+18
352+18
m/z 549 12F-strictosidine
170+18
144+18
m/z 369 12F-strictosidine
aglycones
144+18
170+18
m/z 371 12F-geissoschizine
144+16
352+16
m/z 547 10-OH-strictosidine
170+16
144+16
m/z 367 10-OH-strictosidine
aglycones
144+16
170+16
m/z 369 11OH-geissoschizine

**Supplementary figure 2. The liquid chromatography (LC) and tandem mass spectrometry (MS/MS) of the alkaloid standards and *in vitro* produced alkaloids used in this study.** (A) The LC profiles of alkaloid standards (left panel) and their MS/MS fragments (right panel). Catharanthine, tabersonine, and secologanin standards (SD) were purchased from Sigma Aldrich (St. Louis, USA). Tryptamine, 7-fluorotryptamine, 5-chlorotryptamine, and 5-hydroxytryptamine were purchased from Cayman Chemicals (Ann Arbor, USA). The purifications and NMR identifications of rest standards were described previously.[1] (B) Producing strictosidine, strictosidine aglycones, and geissoschzine 19-epimers/tetrahydroalstonine using purified, recombinant strictosidine synthase (STR), strictosidine β-glucosidase (SGD), and geissoschizine synthase (GS) from *in vitro* reactions (upper panel) as described previously. [2] Recombinant GS produces both 19*E*, and 19*Z*-geissoschizine epimers as well as small amounts of tetrahydroalstonine by-product. Replacing tryptamine with 7-fluorotryptamine and 5-hydroxytryptamine (serotonin) led to the respective 12-fluorinated and 10-hydroxylated strictosidine, strictosidine aglycones, and geissoschzine/tetrahydroalstonine derivatives (left panel). The MS/MS ions of these alkaloid were increased by 18 (fluorine) and 16 (OH) as expected from the respective substitutions (right panel). The mass fragmentation pattens and the addition of 18/16 *m/z* values were used to identify the fluorinated and hydroxylated catharanthine/tabersonine in strain 6 and 7 (Supplementary fig. 6).

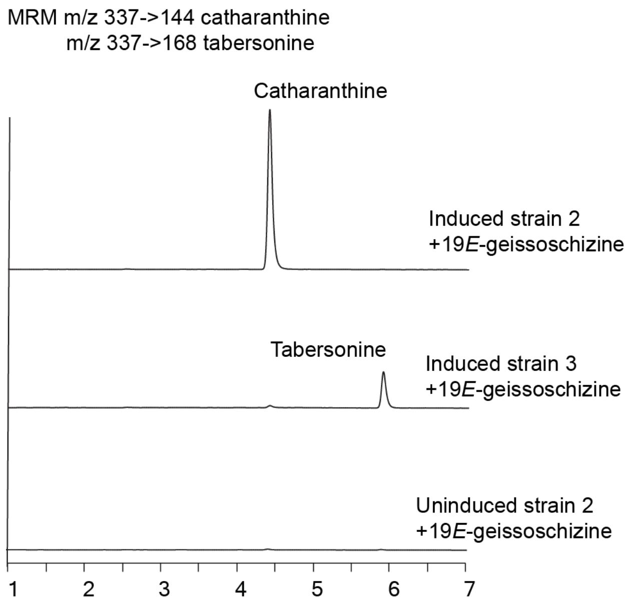


**Supplementary figure 3. Catharanthine and tabersonine were produced in Baker's yeast by feeding 19*E*-geissoschizine to strain 2 containing hydrolase 1, and to strain 3 containing hydrolase 2.** The production of recombinant enzymes was induced by replacing glucose in the media with galactose. The LC-MS/MS analyses showed production of these two alkaloids using the multiple reaction monitoring (MRM) mode. The standards and their MS/MS patterns used for choosing these MRM parameters are found in Supplementary figure 2.

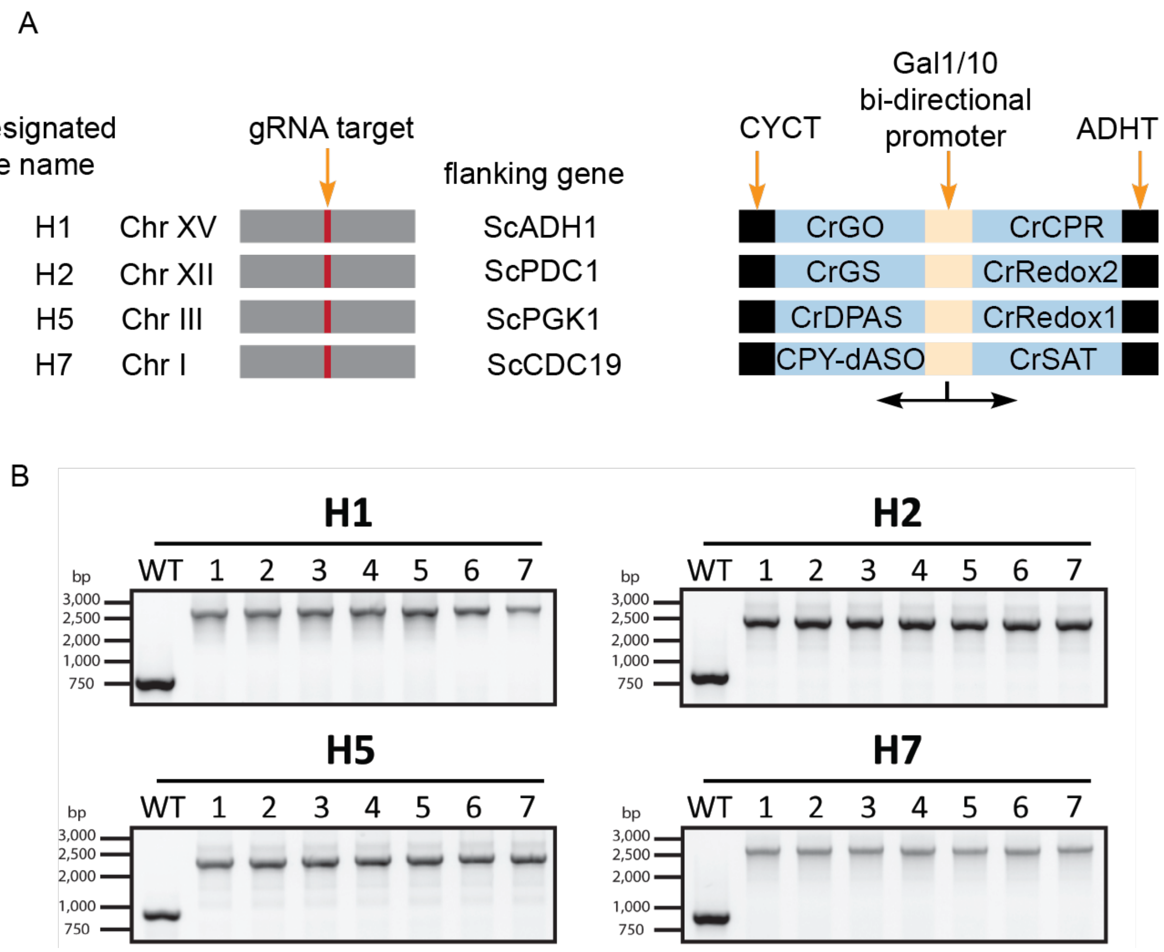


**Supplementary figure 4. Illustration of genomic integrations of 8 alkaloid biosynthetic genes in baker's yeast genome by CRISPR-Cas9 mediated homologous recombination.** (A) A yeast strain (BY4742 MATα, HIS3Δ1, LEU2Δ0, LYS2Δ0, URA3Δ0, designated as SBY104) was previously constructed to contain 4 identical gRNA binding sequences at 4 genomic loci designated as H1, H2, H5, and H7 on 4 yeast chromosomes (XV, XII, III, and I), which are flanked by high-expressing yeast genes ADH1, PDC1, PGK1 and CDC19. [3] The 8 alkaloid biosynthetic genes previously cloned in pESC-Leu, His, and Ura vectors under the control of the Gal1/10 bi-directional promoters, which also included transcription terminators from yeast genes CYC (CYCT) and ADH (ADHT). The respective cassettes were subcloned in pTarget-H1, H2, H5, and H7 vectors and integrated to yeast genome as single copies as described. [3] (B) The integration of respective expression cassette was verified by genomic PCR using 7 independent colonies. Three primers, which would produce an 800 bp product for negative colonies (WT SB104 control lane) and a 2-3 kb product for positive colonies, were used to genotype each colony in each integration site. The primers for genotyping are listed in Supplementary Table 1 with loci H1 using primer 41, 49, 53; H2 using primer 43, 50, 54; H5 using primer 45,51,55; and H7 using primer 47,52,56 (Supplementary table 1).

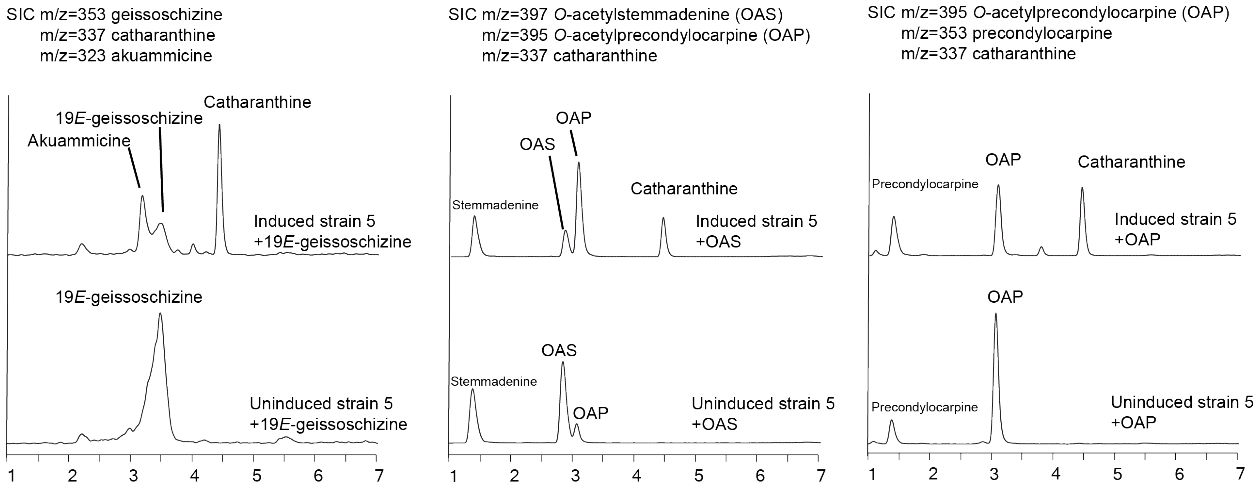


**Supplementary figure 5. Feeding the substrates 19*E*-geissoschizine, *O*-acetylstemmadenine (OAS) and *O*-acetylprecondylocarpine (OAP) led to accumulations of catharanthine and other alkaloids in yeast strain 5.** Three alkaloid substrates were fed to yeast strain 5 after enzyme induction by galactose. OAS and OAP contain labile acetate groups and their deacetylation was observed in uninduced yeast, resulting stemmadenine when compared to the standard (Supplementary figure 3) and precondylocarpine, which was tentatively identified by *m/z* values. OAS slowly but spontaneously oxidizes to OAP at ambient temperature, which was evident in its spontaneous conversion to OAP in the uninduced yeast control. Various amounts of catharanthine were formed in these feeding experiments.

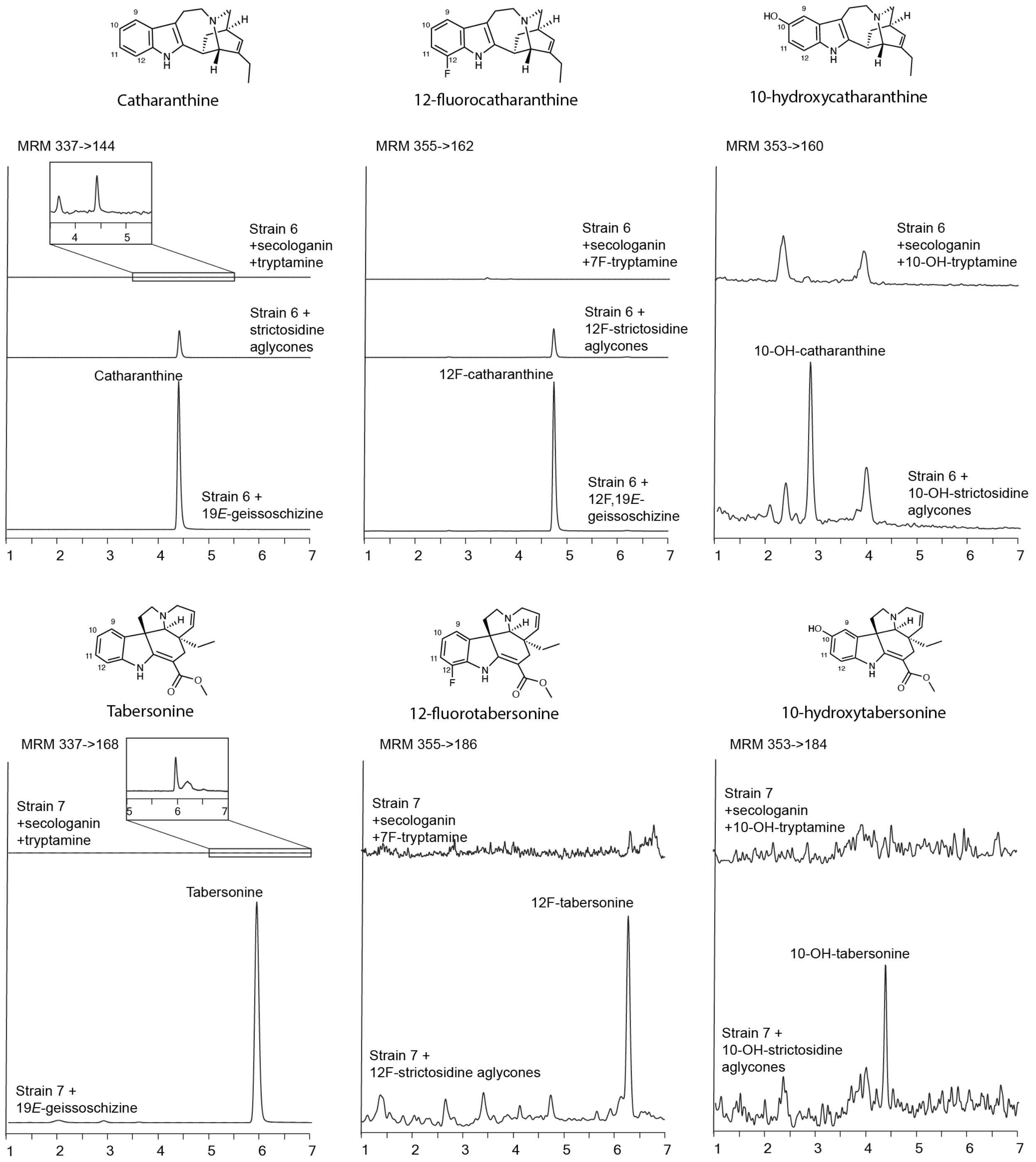


**Supplementary figure 6. Catharanthine, tabersonine and their 12-fluorinated or 10-hydroxylated derivatives were produced by strain 6 and 7.** The aglycones, geissoschizine, and their 12-fluorinated or 10-hydroxylated derivatives were produced *in vitro* as shown in Supplementary fig. 2. The MS/MS patterns for MRM parameters are found in Supplementary fig. 2. The accumulation of catharanthine and tabersonine were detected from secologanin/tryptamine and late intermediates feeding (left panel). The 12-fluorinated catharanthine and tabersonine (middle panel) and the 10-hydroxylated catharanthine and tabersonine (right panel) could be detected from using respective aglycones/geissoschizine derivatives.

### Supplementary table 1. Primers used in this study

| # | Primer name | Primer sequence (5'-3') |
|---|---|---|
| 1 | GO-ApaI-F | TAATGGGCCCATGGAGTTTTCTTTCTCCTCACCAGCTCT |
| 2 | GO-SalI-R | CACCGTCGACATCGTTAACAAGATGAGGAACCAATTT |
| 3 | CPR-NotI-F | AGCGGCCGCATGGATTCTAGCTCGGAGAAG |
| 4 | CPR-NotI-R | AGCGGCCGCTCACCAGACATCTCGGAGATACCTTC |
| 5 | GS-BamHI-F | TACGGGATCCAATGGCTGGAGAAACAACAAAACTAGAC |
| 6 | GS-SalI-R | AGGCGTCGACTCATTCCTCAAATTTCAATGTATTTCC |
| 7 | Redox1-NotI | ATAGCGGCCGCGATGGCTGATCGCGTGAAGA |
| 8 | Redox1-SpeI | ATCACTAGCCAGACAGCTACTGTTGCATT |
| 9 | Redox2-NotI | ATAGCGGCCGCAATGGAAAAGCAAGTTGAGAT |
| 10 | Redox2-SpeI | GTTACTAGTAACTCGAGCAAGTCTCCATCCCAAAGCTC |
| 11 | DPAS-BamHI-F | ATAGGATCCAATGGCCGGAAAATCAGCAGAAGAA |
| 12 | DPAS-SalI-R | ATAGTCGACTTATAACTCTGACGGAGGAGTCAAGGTA |
| 13 | Redox1-BamHI | GTAGGATCCGATGGCTGATCGCGTGAAGA |
| 14 | Redox1-SalI | ATCGTCGACTCAGACAGCTACTGTTGCATT |
| 15 | Gal10-DraIII-F | GTCAATCACTACGTGAATAAGAAGTAATACAAACCGAAAA |
| 16 | ADHT-NaeI-R | GTCAATGCCGGCCAGCTGAATTGGAG |
| 17 | SAT-NotI-F | AGCGGCCGCATGGCACCCCAGATGCAGATA |
| 18 | SAT-SpeI-R | GGACTAGTATTGCTAAAATCAGTGTCCAGA |
| 19 | nativeASO-NotI-F | ATAAGCGGCCGCAATGATAAAAAAAGTCCCAATAGTTC |
| 20 | nativeASO-SpeI-R | ATAACTAGTGGAAGTTCGACTTGTAAATGGAGA |
| 21 | CPY-ASO-BamHI-F | ATAGGATCCGATGAAGGCTTTCACTTCTT |
| 22 | CPY-ASO-SalI-R | ATAGTCGACCAATTCGACTTGCAAGTGT |
| 23 | HL1-NotI-F | ATCGCGGCCGCATGAATTCCTCAACTAATCC |
| 24 | HL1-SpeI-R | CGGCACTAGTTCATGTTTGATGAAAGATGCTA |
| 25 | HL2-NotI-F | ATTAGCGGCCGCATGGGTTCCTCAGATGAGACTATTTTT |
| 26 | HL2-SpeI-R | CGGCACTAGTCTTGATGAAAGAAGCTAAACGTCTG |
| 27 | SAT-SalI-F | ACAAGTCGACATGGCACCCCAGATGCAGATA |
| 28 | SAT-HindIII-R | CTCAAGCTTAATTGCTAAAATCAGTGTCCAGA |
| 29 | SGD-BamHI-F | GAAAGGATCCATGGGATCTAAAGATGATCAGTC |
| 30 | SGD-SalI-R | GAACTCGAGCTTAGTATTTTTGCTTCTTGAC |
| 31 | H1-GO-CPR-F | CATATATTAATTAAAGTCCAATGCTAGTAGAGAAGGCTTCGAGCGTCCCAAAAC |
| 32 | H1-GO-CPR-R | GAAAAGAGCGCGGAGGGGTGTCTGCAGAGCGACCTCATGCTATAC |
| 33 | H2-GS-Redox2-F | GCAGAAACTAACTTCTTCTTCATGTAATAAACACACGGATCCGAGCGACCTCATGCTATACCTGAGAAA |
| 34 | H2-GS-Redox2-R | GATATAAGGTGTTGAAGTTTAGAGATAGGTAAATAAACGCGGATCCCTTCGAGCGTCCCAAAACCTTCT |
| 35 | H5-DPAS-Redox1-F | GGGCGCGAATCCTTTATTTTGGCTTCACCTAGGGAGCGACCTCATGCTATACCTGAGAAA |
| 36 | H5-DPAS-Redox1-R | CTTCCTTTTTCTGGCCCTGATAATAGTATGAGAATTCCTTCGAGCGTCCCAAAACCTTCT |
| 37 | H7-SynASO-F | CTTCACCGGTCGCGTTCCTG |
| 38 | SynASO-SAT-R | ACCGTATCACAACCGACGATCCTTCGAGCGTCCCAAAACCTTC |
| 39 | SynASO-SAT-F | AAAAAAAATAGCCGCCATGACCGAGCGACCTCATGCTATACCTG |
| 40 | H7-SAT-R | CAGGAACGCGACCGGTGAAG |
| 41 | H1 internal primer 5' | GAAGGTGAGACGCGCATAAC |
| 42 | H1 internal primer 3' | GTTTAGCCCATTATGTCTTGTC |
| 43 | H2 internal primer 5' | GTAGCCTCCCCATAACATAAAC |
| 44 | H2 internal primer 3' | CACACGAAAAGTCAGAAGAG |
| 45 | H5 internal primer 5' | GATGACTTCCCATACTGTAATTG |
| 46 | H5 internal primer 3' | GAACCTTCGATTGCTTGTTAC |
| 47 | H7 internal primer 5' | CTGTTCATGGCAACGTCAC |
| 48 | H7 internal primer 3' | GAGCCGAAATTGTGGAAG |
| 49 | H1 CRISPR Confirm | GTAGTGTGCGTGAATGAAGG |
| 50 | H2 CRISPR Confirm | GATTGCAAGGAGAGTGAAAGAG |
| 51 | H5 CRISPR Confirm | AAATCTTGGACAGACAACTTTGAAG |
| 52 | H7 CRISPR Confirm | GGTACCTAGCATCATATGGGAAG |
| 53 | GO-genotyping | CATGGTCCTTTGATGCACGTG |
| 54 | GS-genotyping | GTACTCTGGTGTTTGCAATTTCGATATG |
| 55 | DPAS-genotyping | GTCACACTGATCTTGCCTCAATC |
| 56 | ASO-genotyping | CTGCCTGGGTTGAATCTGGTG |

**Supplementary table 2. Plasmids used in this study and respective primers for plasmid construction.**

| Plasmid | Genes included in plasmids | Cloning Primers* |
|---|---|---|
| 1 | pESC-Leu-GO (ApaI/SalI)- CPR (NotI/NotI) | 1,2 / 3,4 |
| 2 | pESC-His-GS(BamHI/SalI)-Redox1 (NotI/SpeI) | 5,6 / 7,8 |
| 3 | pESC-His-GS(BamHI/SalI)-Redox2 (NotI/SpeI) | 5,6 / 9,10 |
| 4 | pESC-His-DPAS (BamHI/SalI)-Redox1 (NotI/SpeI) | 11,12 / 7,8 |
| 5 | pESC-His-Redox1(BamHI/SalI)-Redox2 (NotI/SpeI) | 13,14 /7,8 |
| 6 | pESC-His-DPAS (BamHI/SalI)-Redox1 (NotI/SpeI)-Gal10-Redox2-ADHT | 15,16 |
| 7 | pESC-Leu-GO (ApaI/SalI)- SAT (NotI/SpeI) | 1,2 / 17,18 |
| 8 | pESC-Leu-GO (ApaI/SalI)- native ASO (NotI/SpeI) | 1,2 / 19,20 |
| 9 | pESC-Ura-CPY-ASO (BamHI/SalI) | 21,22 |
| 10 | pESC-Ura-CPY-dASO (BamHI/SalI) | (Created by SacI digestion) |
| 11 | pESC-Ura-CPY-dASO (BamHI/SalI)-HL1 (NotI/SpeI) | 19,20 / 23,24 |
| 12 | pESC-Ura-CPY-dASO (BamHI/SalI)-HL2 (NotI/SpeI) | 19,20 / 25,26 |
| 13 | pESC-Ura-SAT (XhoI/HindIII)-HL1 (NotI/SpeI) | 27,28 / 21,22 |
| 14 | pESC-Ura-SAT (XhoI/HindIII)-HL2 (NotI/SpeI) | 27,28 / 23,24 |
| 15 | pESC-His-dSTR(NoI/ClaI)-SGD (BamHI/Sal) | Digested from pUG19 / 29,30 |
| 16 | pBluescript-H1-GO-CPR | 31,32 |
| 17 | pBluescript-H2-GS-Redox2 | 33,34 |
| 18 | pBluescript-H5-DPAS-Redox1 | 35,36 |
| 19 | pBluescript-H7-dASO-SAT | 37,38 / 39,40 |

*Cloning primers are listed in Supplementary table 1.

**Supplementary table 3. Yeast backgrounds and plasmid used for transforming each strain.**

| Strain name | Yeast background | Plasmid * |
|---|---|---|
| Strain 0 | BY4742 | 1, 5 |
| Strain 1 | BY4742 | 6, 7, 11 |
| Strain 2 | BY4742 | 6, 7, 12 |
| Strain 3 | BY4742 | 6, 8, 13 |
| Strain 4 | SBY104+GS-GO-CPR-Redox1-Redox2-SAT-CPYdASO-DPAS | 11 |
| Strain 5 | SBY104+GS-GO-CPR-Redox1-Redox2-SAT-CPYdASO-DPAS | 1,11 |
| Strain 6 | SBY104+GS-GO-CPR-Redox1-Redox2-SAT-CPYdASO-DPAS | 1,11,15 |
| Strain 7 | SBY104+GS-GO-CPR-Redox1-Redox2-SAT-CPYdASO-DPAS | 1,12,15 |

*The plasmids are listed in Supplementary table 2.

**References**

1. Qu, Y. *et al.* (2018) Solution of the multistep pathway for assembly of corynanthean, strychnos, iboga, and aspidosperma monoterpenoid indole alkaloids from 19 E-geissoschizine. *Proceedings of the National Academy of Sciences 21*, 201719979–6.

2. Qu, Y. *et al.* (2017) Geissoschizine synthase controls flux in the formation of monoterpenoid indole alkaloids in a Catharanthus roseus mutant. *Planta 25*, 1–10.

3. Baek, S., Utomo, J. C., Lee, J. Y., Dalal, K., Yoon, Y. J., and Ro, D. K. (2021). The yeast platform engineered for synthetic gRNA-landing pads enables multiple gene integrations by a single gRNA/Cas9 system. *Metabolic Engineering 64*, 111–121
